# Zika virus induces persistent phenotypic changes in natural killer cells distinct from dengue virus infection

**DOI:** 10.1101/2025.09.15.676203

**Authors:** Trisha R. Barnard, Noah G. Cruz, Michelle Nguyen, Davis Beltrán, Kassandra Pinedo, Soneida DeLine-Caballero, Uma M. Mangalanathan, Dimelza Araúz, Adriana Weeden, José A. Suarez, Sandra López-Vergès, Catherine A. Blish

**Affiliations:** Department of Medicine, Stanford University School of Medicine, US; Stanford University School of Medicine Postbaccalaureate Experience in Research, Stanford, US; Department of Research in Virology and Biotechnology, Gorgas Memorial Institute for Health Studies (ICGES), Panama City, Panama; Flow Cytometry Core, ICGES, Panama City, Panama; Clinical research and tropical medicine unit, ICGES, Panama City, Panama; Institute for Scientific Research and Technology Services (INDICASAT-AIP), Panama City, Panama; Sistema Nacional de Investigación (SNI-AIP), SENACYT, Panama City, Panama; Chan Zuckerberg Biohub, San Francisco, US

## Abstract

Zika virus (ZIKV) is a mosquito-borne orthoflavivirus of global concern due to its ability to cause congenital neurological defects in infants. Natural killer (NK) cells are innate lymphocytes that can directly kill virus-infected cells. It is thought that NK cells are protective against ZIKV, although the mechanisms by which NK cells detect and respond to ZIKV are not well understood. Here, we evaluated NK cell receptor expression on isolated NK cells from a Panamanian cohort of ZIKV-infected participants, and corresponding NK receptor ligand expression on participant PBMCs and ZIKV-infected cells *in vitro* using mass cytometry. We found that during acute ZIKV infection, NK cells express high levels of activation markers, proliferative markers, and cytotoxic effector proteins, indicating that NK cells are mounting a response to ZIKV. Interestingly, many markers elevated during acute infection remain elevated post-acute infection, suggesting ZIKV infection may have potential long-term effects on NK cell function. Analysis of NK receptor ligand expression on ZIKV-infected participant PBMCs and ZIKV-infected cells *in vitro* did not identify a cellular source of NK cell activation, suggesting that either soluble or tissue-specific factors are responsible for modulating NK cell activity during ZIKV infection. Comparison of the NK cell receptor expression with a previously characterized cohort of dengue-infected participants revealed both common and virus-specific changes in NK cell phenotype during acute infection. This work improves our understanding of the NK cell response to orthoflavivirus infection, which will aid in the development of vaccines and therapeutics.

## Introduction

Orthoflaviviruses, including dengue virus (DENV), yellow fever virus (YFV), and Zika virus (ZIKV), remain a significant public health risk, causing an estimated 400 million infections annually [1]. Infection can cause a range of disease outcomes ranging from asymptomatic or mild fever to severe hemorrhagic or encephalitic disease and death [2]. Specifically, ZIKV remains of global concern due to its ability to cause congenital neurological defects. In healthy adults, the majority of ZIKV infections are asymptomatic or cause a mild febrile illness, suggesting the existence of immune mechanisms mediating protection from disease [3].

Natural killer (NK) cells may be one mediator of disease protection following ZIKV infection, as they are innate lymphocytes that are important effectors in the antiviral immune response. In addition to facilitating immune signaling through the release of cytokines, NK cells can directly kill virus-infected cells through the release of cytotoxic granules containing perforin and granzymes, death receptor-induced apoptosis; and antibody-dependent cellular cytotoxicity. NK cell function is regulated by inhibitory and activating cell surface receptors that recognize ligands on surrounding cells and by cytokines. Consequently, varied expression of activating, inhibitory and cytokine/chemokine receptors on individual NK cells results in phenotypic and functional diversity [4–6]. Virus infection is a known modulator of NK cell function and diversity [7]. NK cells are thought to affect disease severity and contribute to viral control for several related orthoflaviviruses including DENV, YFV, West Nile virus (WNV), and tick-borne encephalitis virus (TBEV), even though the specific mechanism of protection remains unclear [8–13]. Immune profiling of ZIKV-infected participants suggest that NK cells are activated to produce cytokines during acute infection [14]. Yet how ZIKV infection alters different NK cell subsets, how this compares to other related viruses, and whether this affects functional responses remains unclear.

Here, we investigated the NK cell response to ZIKV infection from a Panamanian cohort of ZIKV-infected participants using mass cytometry by time-of-flight (CyTOF). Using two complementary CyTOF panels, we evaluated NK cell receptor expression on isolated NK cells during acute ZIKV infection and after clinical recovery (1-6 weeks post-acute infection). To investigate the potential source of altered NK cell phenotype, we measured corresponding NK receptor ligand expression on ZIKV-infected cells *in vitro* and on patient peripheral blood mononuclear cell (PBMC) samples. Furthermore, we compared NK cell responses in our ZIKV cohort with a previously characterized cohort of acute DENV infection to identify common features and virus-specific aspects of the NK cell response. We found that NK cells are activated during acute ZIKV infection with distinct changes in receptor expression when compared to acute DENV infection. Furthermore, NK cells remain altered for several weeks post ZIKV infection in a manner that is not explained by changes in circulating NK receptor ligand expression.

## Methods

### Zika participant cohort and ethical statement

Febrile patients who attended the Clinical Research and Tropical Medicine Unit at Gorgas Memorial Institute for Health Studies (ICGES) during the 2016-2017 ZIKV outbreak in Panama City, were recruited in the ZIKV participant cohort after informed consent. Sera and blood samples were taken for diagnosis and research, under the protocol approved by ICGES IRB committee (1065/CBI/ICGES/16). Demographic (age, sex), epidemiologic and clinical (days on symptoms onset at recruitment, symptoms and signs) data were collected for each participant. Healthy Panamanian control participants volunteered at Gorgas Memorial Institute of Health Studies and were confirmed to be DENV IgM negative by ELISA (PanBio, Abbott) [15].

### Sera and PBMC Sample Processing & storage

Sera samples were used for diagnosis by specific RT-qPCR for ZIKV, DENV, and Chikungunya virus (CHIKV) [3,16,17]. PBMCs were isolated from EDTA anticoagulated tubes obtained from ZIKV suspected patients; the obtained 12 ml of blood was mixed with 1X Hanks’ Balanced Salt Solution (HBSS) in a ratio of 3:2 (HBSS:anticoagulated blood) and added overlayed on 15 ml of Ficoll-Paque for density gradient centrifugation at 800 × g for 30 min at 20 °C without brake. The PBMC fraction, located at the plasma–Ficoll interface, was carefully collected, washed twice with sterile HBSS (400 × g, 10 min, 20 °C), and resuspended. Cell viability and concentration were determined by trypan blue exclusion using a Neubauer chamber. For cryopreservation, cells were adjusted to 1 × 10^7^ cells/mL in freezing medium (90% fetal bovine serum, 10% DMSO), aliquoted into cryovials, placed in a controlled-rate freezing container (Mr. Frosty), and stored initially at −80 °C for a maximum of 27 h before transfer to liquid nitrogen for long-term storage.

### Mass Cytometry Staining and Data Acquisition

Frozen PBMC samples were thawed in a 37°C water bath and resuspended in 10 mL of RPMI-1640 (Thermo Scientific #21870092) supplemented with 10% fetal bovine serum (FBS, Corning Ref #35-016-CV), 2 mM L-glutamine (Thermo Scientific #25030081), 1% Penicillin/Streptomycin/Amphotericin B (PSA, Gibco #15240062), and 20µL of benzonase (Millipore #70664). One million PBMCs per sample were set aside at 37 °C for staining with the PBMC panel after NK cell isolation (**Table S1**). For samples with greater than 2 million PBMCs, NK cells were isolated using a negative-selection NK Cell isolation kit (Miltenyi Biotec #130-092-657) following the manufacturer’s instructions before CyTOF staining with the NK cell panel (**Table S2**). Samples were stained for mass cytometry as previously described, with the following modifications [18,19]. Live-cell barcoding was performed using anti-β2-microglobulin (clone 2M2; BioLegend #316302) conjugated to Cell-ID cisplatin (^194^Pt, ^195^Pt, ^196^Pt, and ^198^Pt; Fluidigm/Standard BioTools #201194, 201195, 201196, and 201198) and combined in a four-choose-two scheme for 6-plex barcoding [19]. Samples were stained with a (Ethylenediamine) palladium(II) chloride (Sigma-Aldrich #574902, CAS #15020-99-2) viability stain as previously described [19] prior to surface staining with reconstituted lyophilized NK panel or frozen PBMC panel with some Abs spiked-in (**Tables S1-S2**) [15,18,20]. Samples were fixed in 4% PFA in PBS, permeabilized with eBioscence perm buffer (ThermoFisher), and stained with intracellular antibodies prior to storage in an iridium DNA intercalator /2% PFA in PBS (Cell ID Intercalator (^191^Ir, ^193^Ir); Fluidigm/Standard BioTools (#201192B)) at 4°C overnight [19]. Samples were washed once with PBS, three times with water, and mixed with 1x EQ beads (Fluidigm) for normalization. Data was acquired on a Helios mass cytometer.

### Mass Cytometry data pre-processing

Samples were normalized and debarcoded with the premessa R package [21]. EQ beads, doublets, dead cells, and debris were manually gated out in FlowJo v10.10.0 (**Fig S1**). For samples stained with the NK cell panel, any non-NK cells that were not removed during NK isolation were also removed by gating on lineage negative cells (CD3^-^/CD4^-^/CD14^-^/CD19^-^/CD33^-^). Exported FCS files were imported into RStudio using the Flowcore Bioconductor package and converted into SCE format using the CATALYST R package [22,23]. Data was arcsinh transformed with cofactor 5 prior to downstream analysis. Samples with <70% viability at thaw or with fewer than 10,000 (PBMC) or 500 (NK) cells in the final exported fcs file were excluded from the analysis.

### Unsupervised clustering of Mass Cytometry Data

Cell subsets were identified by unsupervised clustering using parc [24]. NK cells were clustered on all markers other than the lineage markers used to exclude non-NK cells and resolution parameter 0.1, chosen because other resolutions did not identify a canonical CD56^bright^CD16^-^ cluster of NK cells. To identify coarse cell types, the PBMC data was clustered on lineage markers (CD19, CD20, CD3, CD4, CD8, CD56, CD7, CD14, CD16, CD11b, CD33, BDCA2, CD163, CD64, CD1C, CD11c, CD141, and HLA-DR) using resolution parameter 0.12. This and all other clustering resolutions of PBMC data identified several subsets of CD4 and CD8 T cells, as well as a HLA-DR-expressing cluster of undetermined cell type that trended toward increased frequency in acute infection (p=0.056, **Fig S4ab**). To identify which cell types comprised this compartment, cells in this cluster were subsetted and re-clustered with resolution parameter 0.12 (**Fig S4c**). For the final cell type identification, clusters 1 and 5 were manually merged into the final CD4 T cell cluster, clusters 2 and 4 were manually merged into the final CD8 T cell cluster, and cluster labels from the subsetted re-clustered undetermined cluster 8 were assigned to the original object. Differential state testing was performed using the diffcyt R package [25].

### CytoGLMM analysis

Use of the CytoGLMM package has been described previously [26]. Only samples with greater than 500 cells per indicated cell type were included in the analysis. Cells were randomly subsampled to an equal amount per donor and all GLM analysis was bootstrapped at least 1000 times. Comparison of Acute and Post-acute infection was performed as a paired analysis; only participants with both an acute and post-acute sample were included.

### Integration with DENV data

We had additional PBMC vials available from some healthy controls analyzed in our previous DENV studies with similar CyTOF panels [15,20], which we analyzed here as part of our healthy control cohort. FCS files from [15] were normalized together with data collected herein using premessa [21]. To directly compare ZIKV and DENV NK cells samples, we used cyCombine to integrate the two datasets with overlapping healthy control samples used as anchor samples [27]. Distributions of all markers on both batches were verified to be similar after integration (**Fig S6**).

### Cell culture and Virus info

K562 cells (ATCC #CCL-243) were maintained in RPMI-1640 supplemented with 10% FBS, 2 mM L-glutamine, 1% Non-essential amino acids (NEAA, Thermo Scientific #11140050), 1 mM Sodium pyruvate (Thermo Scientific #11360070), 10 mM HEPES (Thermo Scientific #15630080), and 1% PSA. A549 (ATCC #CCL-185) and Vero cells (ATCC #CCL-81) were maintained in DMEM (Thermo Scientific #11885092) supplemented with 10% FBS, 2 mM L-glutamine, 1% NEAA, 1 mM Sodium pyruvate, 10 mM HEPES, and 1% PSA. JEG-3 cells (provided by Virginia Winn) were maintained in EMEM (Corning #10009CV) supplemented with 10% FBS, 2 mM L-glutamine, 1% NEAA, 1 mM Sodium pyruvate, 10 mM HEPES, and 1% PSA. ZIKV isolate PRVABC59 (Genbank accession KU501215, obtained through BEI Resources, NIAID, NIH, NR-50240) [28] was propagated in C6/36 cells (ATCC #CRL-1660) followed by one passage in Vero cells. Viral stocks were titered by TCID50 assay; titers were converted to PFU/mL for Multiplicity of Infection (MOI) calculations using 0.7 PFU ≅1 TCID50 [29]. All infections were performed using media containing 2% FBS.

### NK ligand expression on infected cells

Ligand expression was evaluated 24 h post-infection with MOI 1. Adherent cells were dissociated with TrypLE Express (Thermo Fisher #12-604-021) prior to CyTOF or flow cytometry staining. Cells were stained with the CyTOF ligand panel (**Table S1**) without the additional spike-in antibodies as described above for PBMCs, except K562 cells were not barcoded due to low expression of β2-microglobulin.

After collecting CyTOF data from our participant samples, we noticed that the HLA-Bw4 signal from our frozen CyTOF antibody panel was lower than expected on our positive control samples and therefore excluded this marker from CyTOF analyses. We therefore measured HLA-Bw4 expression on ZIKV-infected cells by flow cytometry using clone REA274 (Miltenyi Biotec #130-133-385) and Flavivirus group antigen (clone 4G2, Novus Biologicals #NBP2-52709AF647) to detect infected cells. 4G2 signal was used to identify virus-infected vs. bystander cells in the same culture (**Fig S1**). Mean signal intensities of arcsinh-transformed CyTOF data or geometric Mean Fluorescence intensities of flow cytometry data were scaled individually for each marker for comparison on the heatmap.

### Data visualization and Statistical analysis

Data visualization was performed in R using the ggplot2, ComplexHeatmap, CATALYST, CytoGLMM and cyCombine packages [23,26,27,30,31]. All statistical analysis was performed in R. Unless otherwise noted, differences between Healthy and Infected groups were compared using Wilcoxon rank-sum test with Holm’s correction for multiple comparisons using the ggpubr package [32].

## Results

### NK cells are activated during acute ZIKV infection and remain altered for at least 6 weeks

To understand how NK cells respond to ZIKV infection, we profiled NK cell receptor and ligand expression by mass cytometry [18] on PBMCs from a Panamanian cohort of adult ZIKV participants during the acute phase of infection (1-5 days post symptom onset) and post-acute infection (early and late convalescence, 1-6 weeks later) (**Table 1:** Participant demographics and **Table S2**). We found that CD38, CD69, Fas Ligand, perforin, Ki67, and FcRγ expression was upregulated on NK cells during acute ZIKV infection, suggesting NK cells are activated, proliferating, and potentially cytotoxic during acute infection (**Fig S1**). Additionally, though the activation markers CD38 and CD69 returned to baseline 2-6 weeks post-symptom onset, other markers including perforin, Ki67, and FcRγ remain elevated, suggesting that ZIKV may induce long-lasting (for at least 6 weeks after acute phase) changes in NK cell phenotype.

**Table 1:** Participant characteristics

| Parameter | Acute (n = 11) | Post-acute (n = 9) | Healthy (n = 14) |
| --- | --- | --- | --- |
| Age, y, median (range) | 33 (22-72) | 37 (22-72) | 32 (16-55) |
| Females | 6 | 4 | 9 |
| Males | 5 | 5 | 5 |
| Days post symptom onset, median (range) | 3 (1-5) | 18 (7-49) | NA |

To identify which NK cell subsets were changing during infection, we performed unsupervised clustering on live NK cells (**Fig 1a-b**). This identified 9 clusters, including the canonical CD56^bright^CD16^-^ immature NK cells (cluster 8, 5.52%), adaptive-like CD57^+^NKG2C^hi^ NK cells (cluster 1, 11.7%), proliferating Ki67^hi^ NK cells (cluster 5, 0.03%), and DNAM1^+^ cells with low expression of other NK markers (cluster 4, 0.57%). CD56^dim^CD16^+^ NK cells further separated into 5 additional clusters based on KIR status and Syk expression: KIR^low^Syk^-^ (cluster 3, 6.93%), KIR^low^ Syk^+^ (cluster 7, 39.1%), KIR2DL5^+^ (cluster 9, 0.43%), KIR3DL1^+^ (cluster 2, 11.4%), and KIR2DS4^+^ (cluster 6, 24.3%). Differential abundance analysis revealed a trend toward decreased frequency of CD56^bright^CD16^-^ NK cells during acute ZIKV infection (p = 0.051), suggestive of increased NK activation during acute ZIKV. There were no other changes in cluster frequencies, suggesting that the ZIKV-associated changes in activation/cytolytic markers occurred across multiple NK cell subsets rather than changing the distribution of NK cell subsets (**Fig 1c**). Indeed, during acute infection CD38 expression increases on CD56^bright^CD16^-^ (immature), CD56^dim^CD57^+^ (mature), and CD57^+^NKG2C^hi^ (adaptive) as well as KIR^+^ (educated) and KIR^low^ (uneducated), and CD69 expression increases on all subsets except CD56^bright^CD16^-^ (**Fig S3**). Similarly, perforin and Ki67 expression increased similarly on immature, mature, educated, and uneducated NK cells in acute infection. Interestingly, both Fas Ligand and FcRγ increase only on immature/uneducated NK cells during acute infection, whereas post-acute infection FcRγ is increased on all subsets except for adaptive NK cells which do not express FcRγ (**Fig S3**)[33].

**Figure 1:**
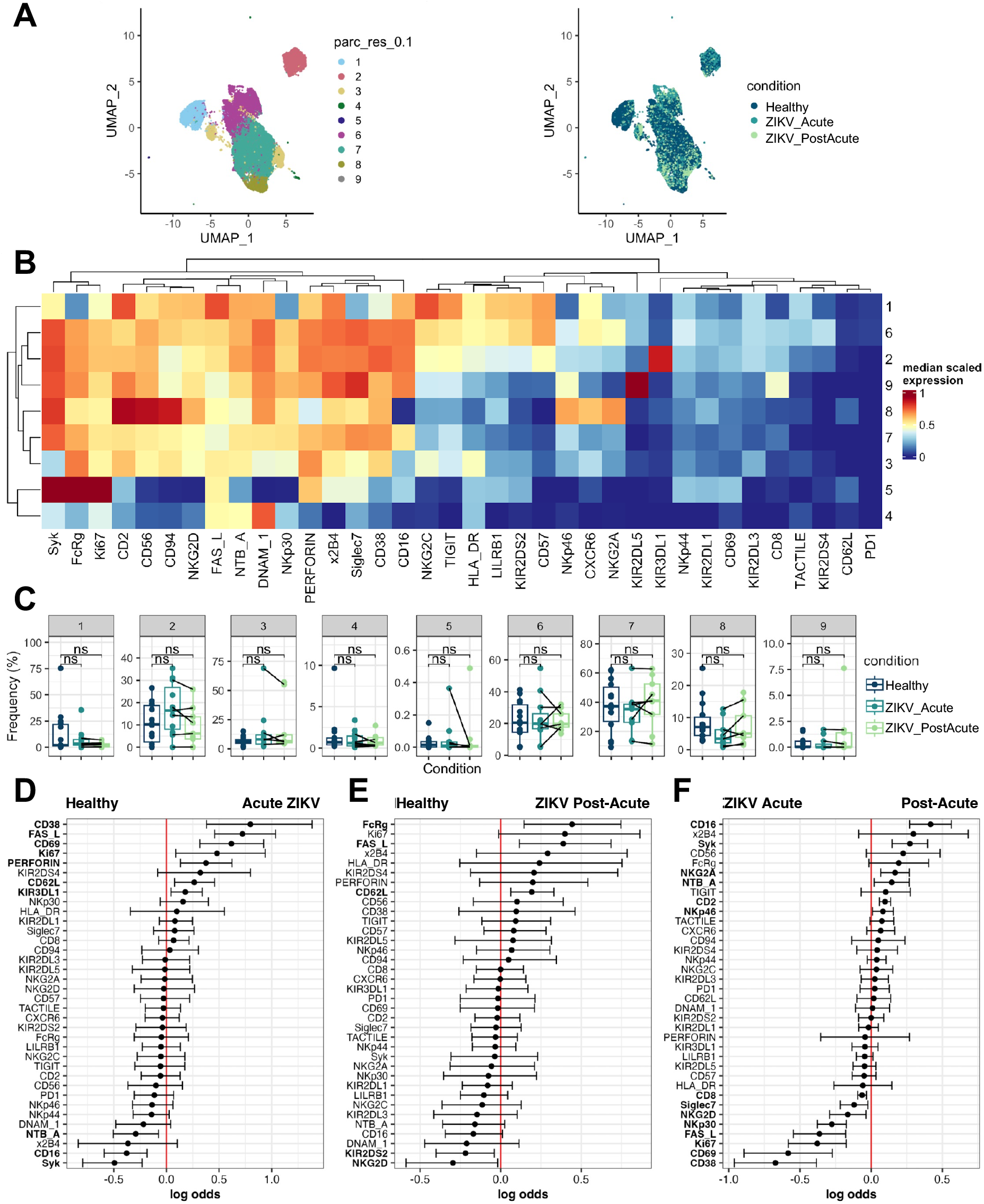
NK cells are activated during acute ZIKV infection and remain altered for at least 6 weeks. (a) UMAP of NK cells colored by cluster and condition (b) Heatmap of median marker expression in each cluster (c) Abundance of each cluster (d) cytoGLM analysis of NK cells from Healthy (n = 13) vs. Acute ZIKV (n = 11) (e) cytoGLM analysis of NK cells from Healthy (n = 13) vs. ZIKV Post-acute (n = 9) (f) cytoGLMM analysis of NK cells from Acute ZIKV vs. ZIKV Post-acute (n = 6 pairs)

Finally, to identify specific NK cell markers that were predictive of ZIKV infection, we used CytoGLM, which uses a generalized linear model to identify which protein expression levels are most predictive of a particular condition [26]. CytoGLM accounts for both person-to-person variability and correlations between proteins [26]. We first evaluated markers predictive of acute ZIKV infection compared to healthy samples, which confirmed elevation of CD38, Fas Ligand, CD69, Ki67, and perforin as observed in our analysis of mean signal intensity (**Fig 1d**). In addition, CytoGLM identified increased KIR3DL1 and CD62L expression as predictive of acute ZIKV infection, and increased NTB-A, CD16, and Syk expression as predictive of healthy status. When comparing healthy to post-acute ZIKV infection, CytoGLM identified FcRγ, Fas Ligand, and CD62L as predictive of post-acute infection when compared to healthy. All of these are markers that were identified as increased during acute infection by either MSI or CytoGLM, indicating potential long-term changes in NK cells that initiate during acute infection (**Fig 1e**). In a paired analysis using the same ZIKV-infected participants during acute and post-acute infection, we identified high CD38, CD69, Ki67, and Fas Ligand expression, and low CD16, Syk, and NTB-A expression as predictive of acute infection (**Fig 1f**). The increased power of the paired analysis also identified high expression of NK activating receptors NKp30 and NKG2D as predictive of acute infection. Finally, FcRγ, perforin and CD62L, markers whose expression is either upregulated or predictive of both acute and post-acute conditions, are not predicted to be different in acute vs. post-acute infection. Overall, the data support that peripheral NK cells are activated, proliferating, and potentially cytotoxic during acute ZIKV infection, and that not all changes in the NK cell compartment return to baseline by 1-6 weeks post-acute infection.

### NK ligand expression on infected cells

To understand how ZIKV infection may alter susceptibility to NK cell killing of infected cells and determine whether ZIKV-infected cells may directly contribute to NK cell activation, we profiled NK cell receptor ligand expression on ZIKV-infected cells *in vitro*. As we wanted to identify cell type-independent changes driven by virus replication itself, we compared the change in expression on infected vs. bystander cells in the same culture for three different cell lines: A549 (lung epithelium carcinoma commonly used to study ZIKV), JEG3 (placental choriocarcinoma, to model ZIKV infection of the placenta), and K562 (myeloid leukemia, to model ZIKV infection of myeloid cells and commonly used to study NK responses) (**Fig 2**) [34–37]. We found that Death receptors (DR4-DR5, TRAIL receptors) decreased only on ZIKV-infected cells on all three cell types, and expression of the NKG2D activating ligands ULBP1-ULBP2 decreased slightly on infected A549 and K562 cells. Class I HLA molecules increased only on bystander A549 cells, but not on other cell types or on infected cells. These results suggest that ZIKV-infected cells may escape NK cell killing via TRAIL and/or NKG2D, but otherwise ZIKV does not drastically alter the NK cell ligand repertoire on infected cells.

**Figure 2:**
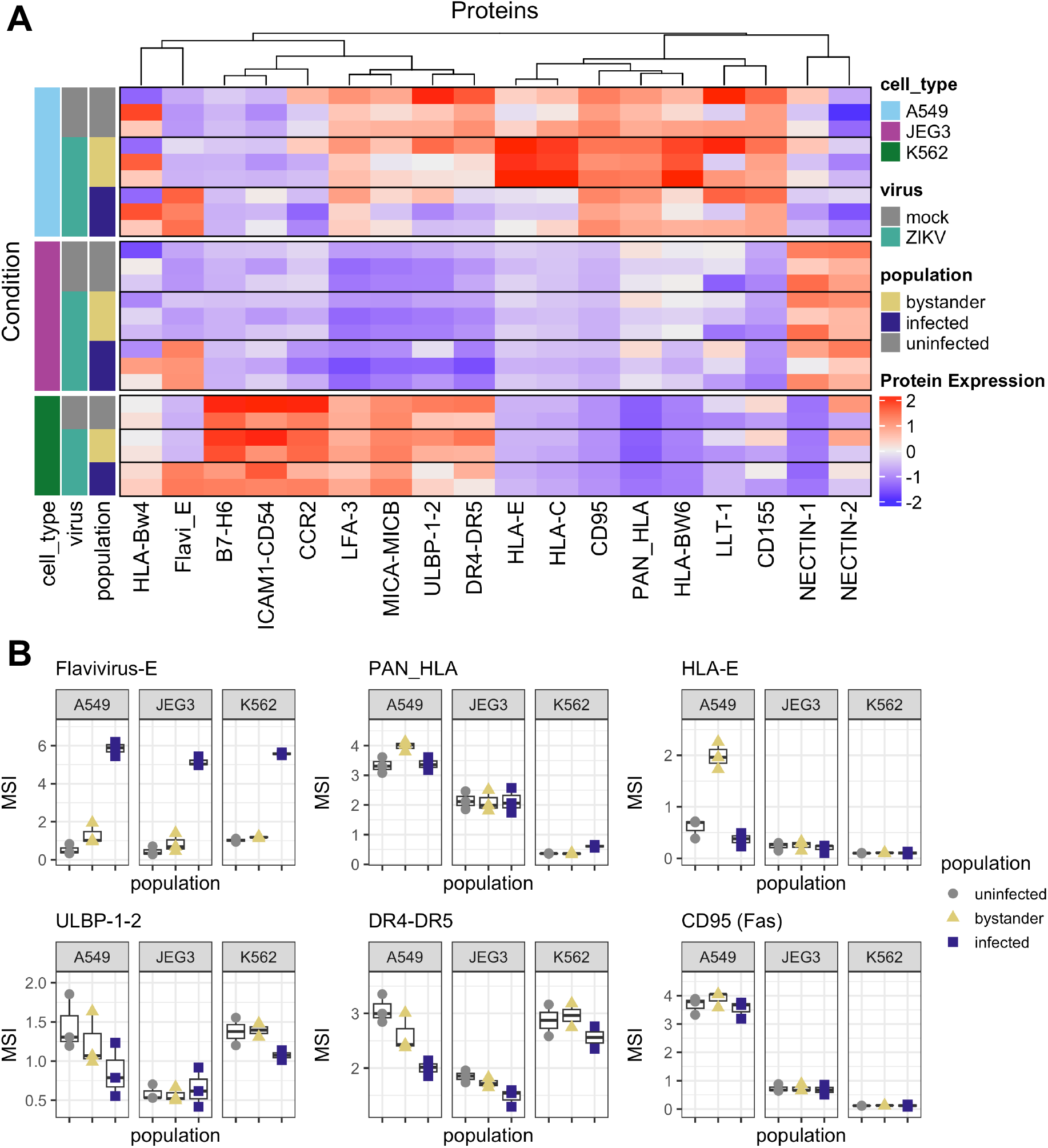
NK receptor ligand expression in ZIKV infected cells. a) Heatmap of mean protein expression in the indicated condition b) Boxplots of mean signal intensity (MSI) of selected proteins

### Altered expression of NK ligands on PBMCs during acute ZIKV infection

To place NK receptor expression in greater context, we analyzed NK ligand expression on ZIKV patient PBMCs by mass cytometry (**Table S1**). We first identified major PBMC cell types using unsupervised clustering at a coarse resolution on lineage markers (**Fig 3a-b and S4**). We identified CD4^+^ T cells (41.4%), CD8^+^ T cells (29.7%), NK cells (18.1%), CD14^+^ myeloid cells (6.4%), CD19^+^CD20^+^ B cells (3.3%), CD19^+^CD20^-^ B cells (0.39%), as well as several additional myeloid cell subsets which included multiple dendritic cell types (0.74% total). Differential abundance analysis showed no significant difference in cluster abundances during ZIKV infection for all clusters except CD19^+^CD20^-^ B cells which expanded in frequency during acute infection (**Fig 3c**).

**Figure 3:**
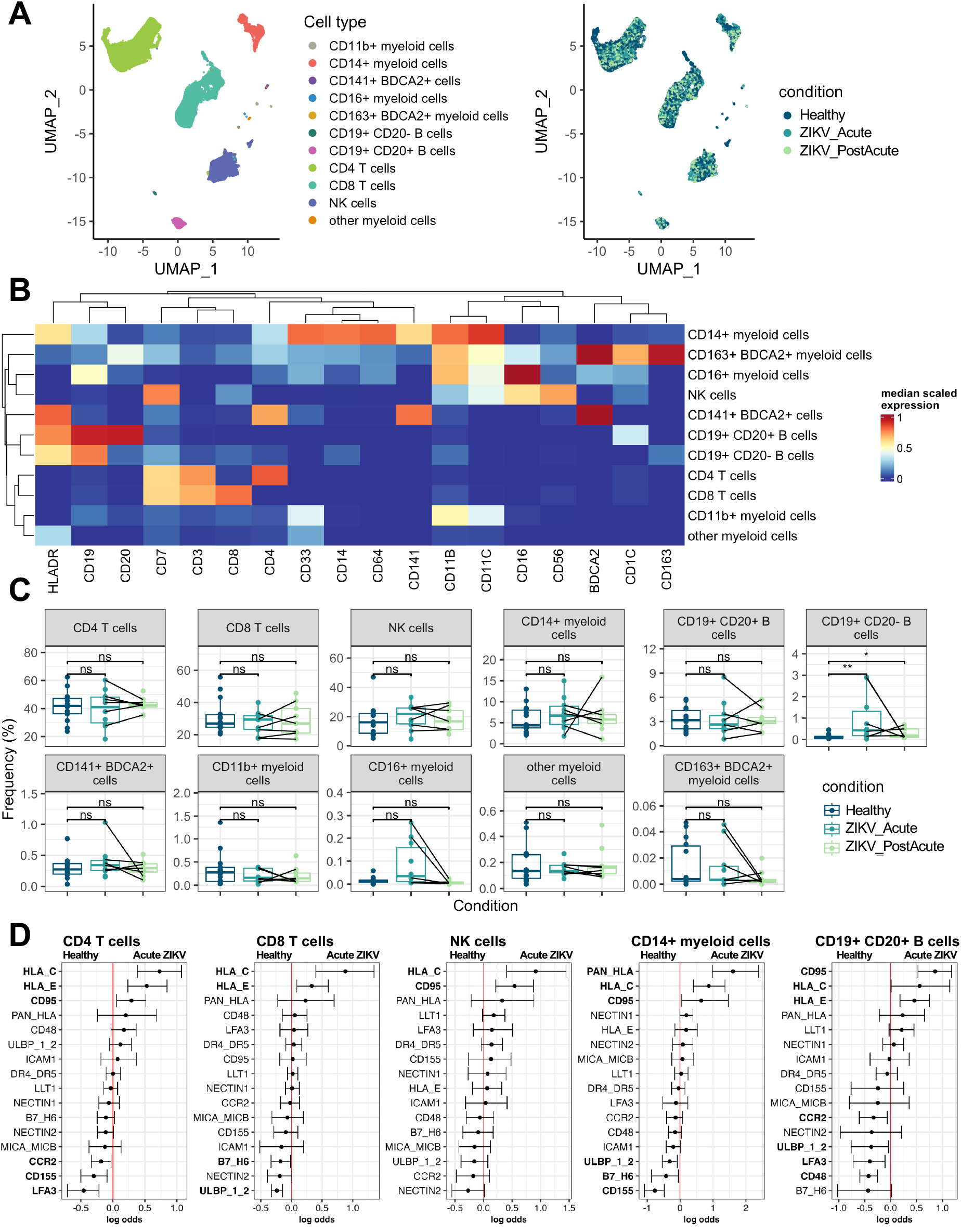
NK receptor ligand expression in major PBMC subsets. a) UMAP of PBMCs colored by cell type and condition b) Heatmap of median marker expression in each cluster c) Abundance of each cluster d) CytoGLM analysis of the indicated cell type in Healthy (n = 13-14) vs. Acute ZIKV (n = 9-10)

To determine how ligands for NK cell receptors are changing on the immune cell compartment during acute and post-acute ZIKV infection, we performed CytoGLM analysis of major PBMC subsets (**Fig 3d and S5**). We found that increased expression of NK inhibitory receptors HLA class I (Pan-HLA; HLA-C, HLA-E) and CD95 (Fas) are predictive of acute ZIKV infection vs. healthy across multiple cell types. Interestingly, unlike in NK cells, most of the proteins that change during acute infection resolve by 1-6 weeks post-acute infection (**Fig S5**). This is supported by differential state testing of identified cluster-marker combinations, which identified 65 significantly altered cluster-marker combinations in acute infection and only one cluster-marker combination significantly altered in post-acute infection compared to healthy (**File S1**). Overall, these data did not identify specific ligands upregulated in PMBCs that accounted for NK cell activation seen in acute infection.

### Comparison with DENV NK response

We previously profiled the NK cell response to mild DENV infection using the same mass cytometry panel [15,20]. To compare the NK cell receptor expression in ZIKV, we used cyCombine to integrate the two datasets [27]. After integration, there were no major differences in marker expression across batches (**Fig 4a and S6**). Overall, NK cell receptor expression was similarly perturbed during acute infection with both viruses. CD38, CD69, Fas Ligand, Ki67, and perforin increased in expression, and Syk, 2B4, and CD56 decreased in expression relative to healthy controls, albeit to different extents in ZIKV vs. DENV. Virus-specific changes in NK cell receptors were an increase in FcRγ expression unique to ZIKV, and increased HLA-DR expression and decreased NKp46, Siglec7, and CD94 expression unique to DENV (**Fig 4b-c**). CytoGLM analysis comparing ZIKV to DENV further identified increased expression of NKG2A, KIR2DL3 and KIR3DL1 as predictive of ZIKV, and increased expression of NKG2D as predictive of DENV. Overall, the NK cell response to ZIKV is similar to the NK response to mild DENV, albeit with slightly different magnitude and overall more perturbations in DENV than ZIKV.

**Figure 4:**
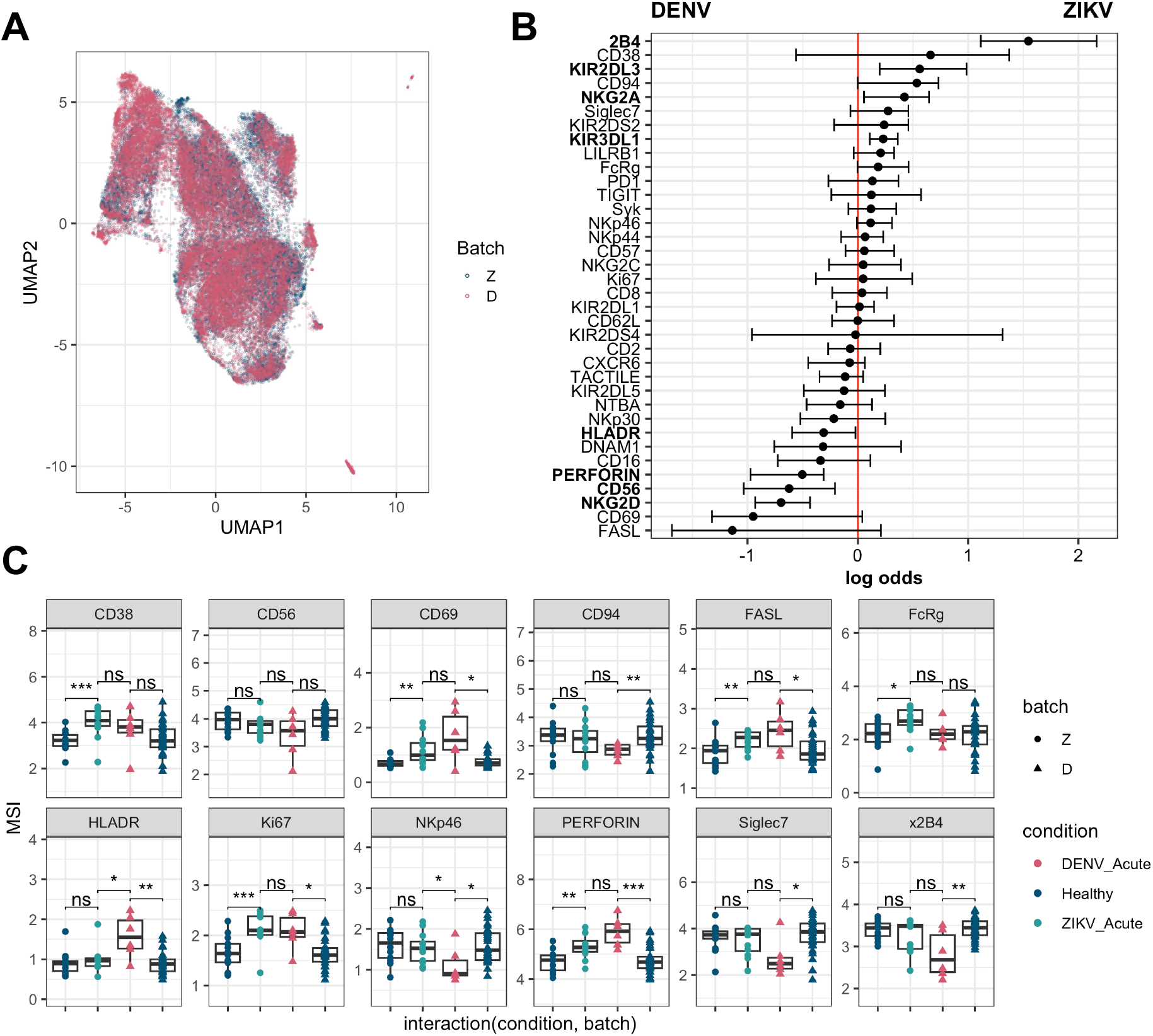
Comparison with DENV profiling. a) UMAP of NK cells colored by batch (Z = this study, D=McKechnie *et al* 2020 [20]) after integration of datasets b) CytoGLM analysis of NK cells in Acute DENV (n = 5) vs. Acute ZIKV (n = 10) c) Mean signal intensities (MSI) of selected protein expression on NK cells

## Discussion

We investigated how ZIKV infection alters the NK cell receptor-ligand repertoire during acute and post-acute infection. We found activation of NK cells during acute infection and elevated perforin, Ki67, and FcRγ expression that lasted for at least 6 weeks post-acute infection. Examination of NK receptor ligand expression on participant PBMCs or on ZIKV-infected cells *in vitro* did not identify a cellular source of persistent NK cell perturbations. Unlike what has been observed in other orthoflavivirus infections, we observed increased expression of NK cell activation and proliferation markers independent of stage of differentiation [8,9,38]. Comparison of the NK cell receptor repertoire with DENV revealed both common and virus-specific changes in NK cell phenotype during acute infection. Our data is consistent with previous studies of acute ZIKV infection that demonstrate modest systemic immune activation during acute ZIKV infection and raise the question of potential long-term effects of ZIKV infection on NK cell function [39].

Previous studies have reported conflicting results as to whether orthoflavivirus infection can lead to long-lasting changes in NK cells. NK cells from people with prior symptomatic West Nile Virus (WNV) infection display an altered phenotype and enhanced IFNγ production in response to *in vitro* WNV infection [10]. Additionally, DENV infection may induce signatures suggestive of activation and exhaustion on adaptive-like NK cells that persists in convalescence [11]. Unlike reports from TBEV and live attenuated YFV 17D, which observe Ki67^+^ NK cells return to normal by 2-3 weeks post-infection, we observe elevated NK cell proliferation for at least 6 weeks post ZIKV infection [8,9]. Interestingly, though we identify persistent changes in NK cell phenotype after ZIKV infection, we do not see evidence that adaptive NK cells expand in response to ZIKV. Rather, ZIKV infection uniquely leads to elevation of FcRγ which is downregulated in adaptive NK cells [40]. FcRγ (also known as FcεRIγ) is an intracellular signaling adaptor that associates with CD3ζ to facilitate signaling from CD16, NKp30, and NKp46 activating receptors [33,41]. Loss of FcRγ is associated with cytomegalovirus (CMV) infection, so elevated FcRγ may be due to reduced CMV prevalence in our participant cohorts [40]. Alternatively, FcRγ expression is associated with NK cell proliferation in response to cytokines [41], which could be driven by IL-2, IL-12, and IL-18, all of which are upregulated in acute ZIKV infection [42,43]. Interestingly, during acute infection FcRγ expression increases the most on immature NK cells which are the most responsive to cytokines [44], whereas in post-acute infection FcRγ increases the most on mature NK cells. It is tempting to speculate that FcRγ expression is initially elevated in immature NK cells and remains elevated while NK cells mature. Additional studies are needed to determine the cause and the consequence of elevated FcRγ in ZIKV infection, and whether NK cells from post-acute ZIKV patients mount improved responses to ZIKV-infected cells or other infectious or non-infectious stimuli using positive receptors associated with FcRγ.

Our data did not identify specific ligands upregulated in PMBCs or infected cells that accounted for the NK cell activation seen in acute infection. The fact that all NK cell subsets are displaying an activation signature in acute infection despite receiving strong inhibitory signals from PBMCs suggests that NK cells may be activated primarily by soluble factors or tissue-specific signals. Indeed, cytokine profiling of ZIKV patients has demonstrated persistent elevation of cytokines known to promote NK cell activation and proliferation [39,42,43]. Interestingly, unlike the NK cells themselves, there did not appear to be lasting changes in expression of NK cell ligands within the other cell types within the PBMC compartment, providing further support for the idea that circulating PBMCs are not responsible for modulating NK cell activation in ZIKV. Additionally, though we identify only minor changes in NK ligand expression on ZIKV-infected cells, it is possible that ligands we did not measure contribute to NK recognition of infected cells [45]. During acute ZIKV infection, circulating PMBCs had increased expression of HLA-C and HLA-E, suggesting a potential for NK cell inhibition, similar to what we found in acute DENV [15]. However, ZIKV-infected cells *in vitro* did not upregulate HLA-E as DENV infection does, suggesting that ZIKV may not inhibit NK cell response against infected cells [46]. Increased expression of Fas Ligand on NK cells suggests that NK cells may attempt to eliminate infected cells via induced apoptosis, though infected cells likely avoid extrinsic apoptosis by reducing signaling downstream of Fas receptor and downregulating the expression of TRAIL receptors, though this may be cell-type dependent [47,48].

This study has several limitations. First, we were unable to reliably detect infected cells in PBMC samples from our cohort, so we are unable to compare NK ligand expression on infected cells *in vivo*. Second, the time post-infection of our post-acute samples varied from 1-6 weeks following the acute infection sample. Therefore, it is possible that though we detected changes in NK cells that lasted beyond acute infection, the NK cell compartment will eventually return to baseline. Additionally, though we don’t have detailed cytokine information in our cohort, cytokine changes in ZIKV infection have been extensively characterized elsewhere (reviewed in [49]). Finally, we were not able to analyze NK cell functions in ZIKV participant samples due to limited sample availability.

To the best of our knowledge, this work represents the most detailed examination of NK cell phenotype in ZIKV infection to date. Though we were unable to identify a cellular source of changes in the NK cells, our profiling of NK ligand expression on ZIKV-infected cells adds to previous work examining NK cell response to ZIKV infected cells. Future work is required to determine the duration of the ZIKV-induced perturbations in the NK cell compartment, as well as whether these changes affect NK cell functions during acute infection into recovery as well as in future infections by similar viruses like DENV or other orthoflaviviruses.

## Data availability

Normalized and debarcoded.fcs files for all mass cytometry data generated in this study along with their accompanying metadata will be deposited at CytoBank Community upon publication. Code used for data analysis will be made available at https://github.com/BlishLab/Zika_NK upon publication.

## Acknowledgements

This work was supported by a Chan Zuckerberg Investigator Award, NIH R21 AI135287 and R21 AI130523 (to CAB) and grants 9044.51 and 3.04.18 from the Ministry of Economy and Finance of Panama and grants SNI 021-2020 and SNI 052-2023 (to SLV). TRB is supported by a fellowship from the Stanford Medicine Pandemic Preparedness Hub. The Stanford Medicine Postbaccalaureate Experience in Research Program and Stanford Medicine’s Division of Infectious Diseases & Geographic Medicine Merigan Scholar Fund supports NGC. MN is supported by a Stanford University Major Grant. We are grateful to members of the Blish lab for helpful comments and Mark M. Davis and the Stanford Human Immune Monitoring Core for use of their mass cytometers. We thank all health institutions, participants, and volunteers for their participation in the study.

## Author Contributions

Conceptualization: TRB DB SLV CAB

Methodology: TRB NGC

Investigation: TRB NGC MN DB KP SDC UMM DA AW JAS

Formal analysis and Visualization: TRB

Writing - Original Draft: TRB

Writing - Review & Editing: TRB NGC DB SLV CAB

Supervision and Funding acquisition: SLV CAB

## Supplemental information

**Fig S1:**
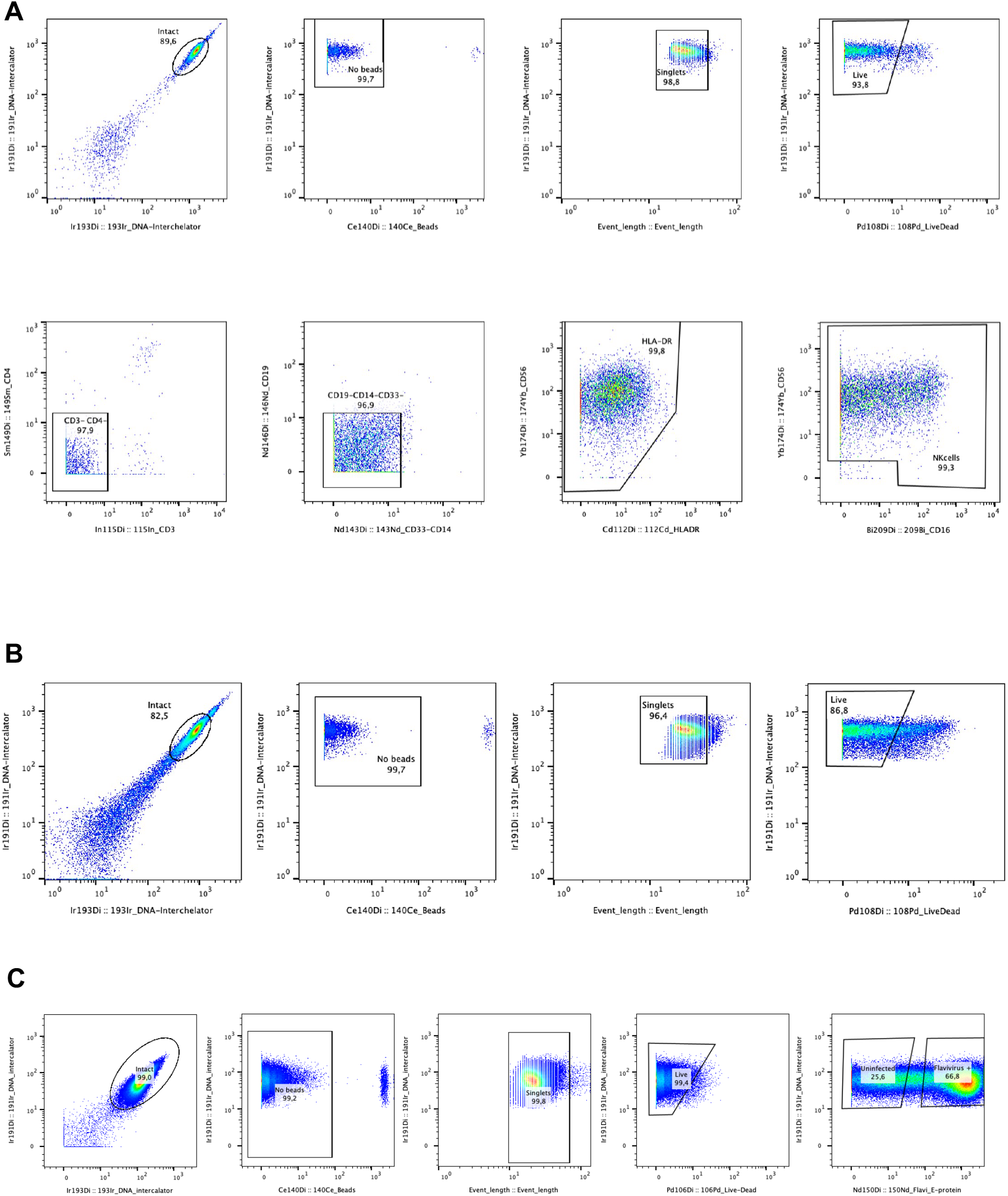
CyTOF gating strategy a) NK cells b) PBMCs c) Infected cells

**Fig S2:**
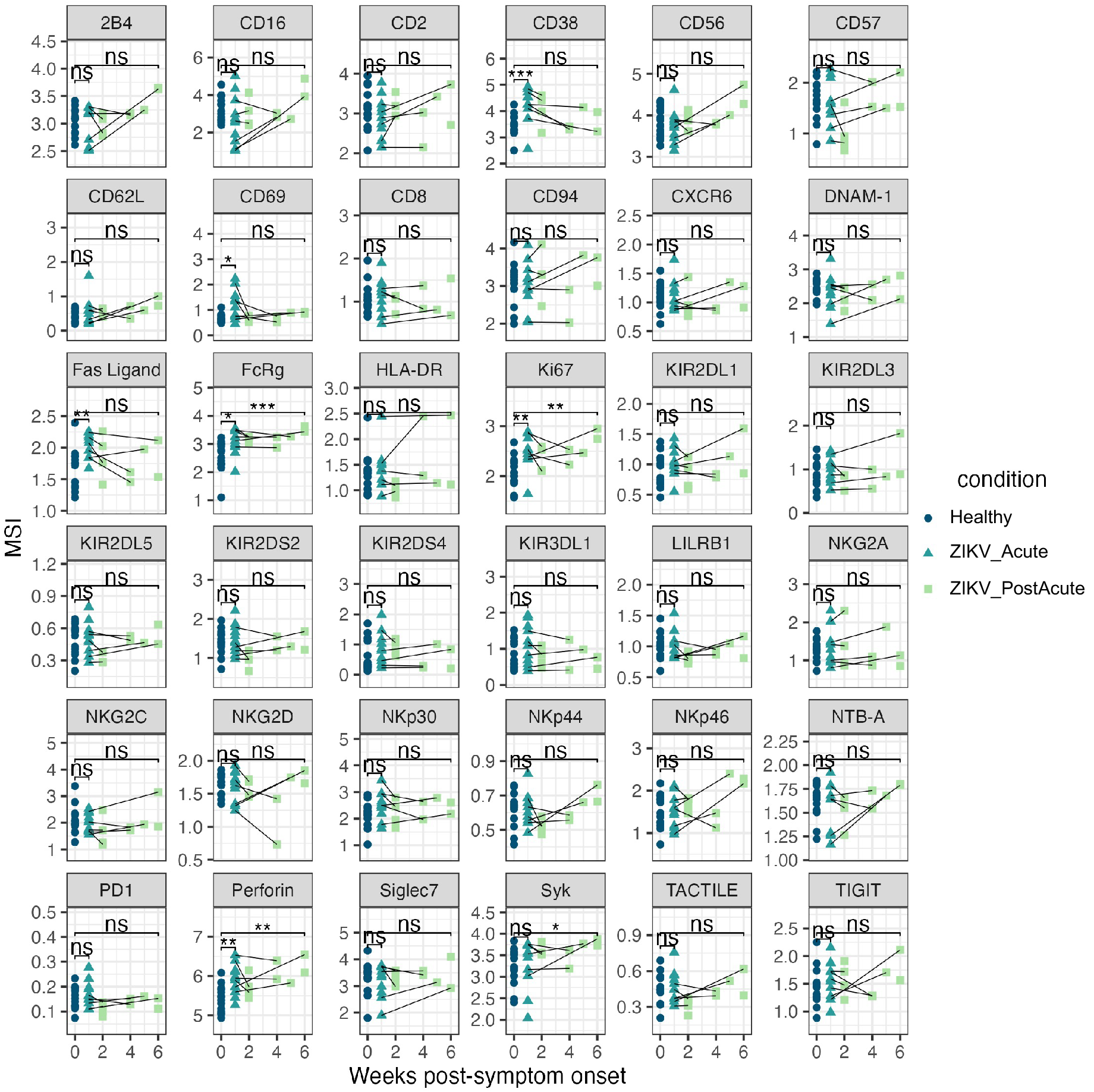
MSI of all NK markers by time post-infection

**Fig S3:**
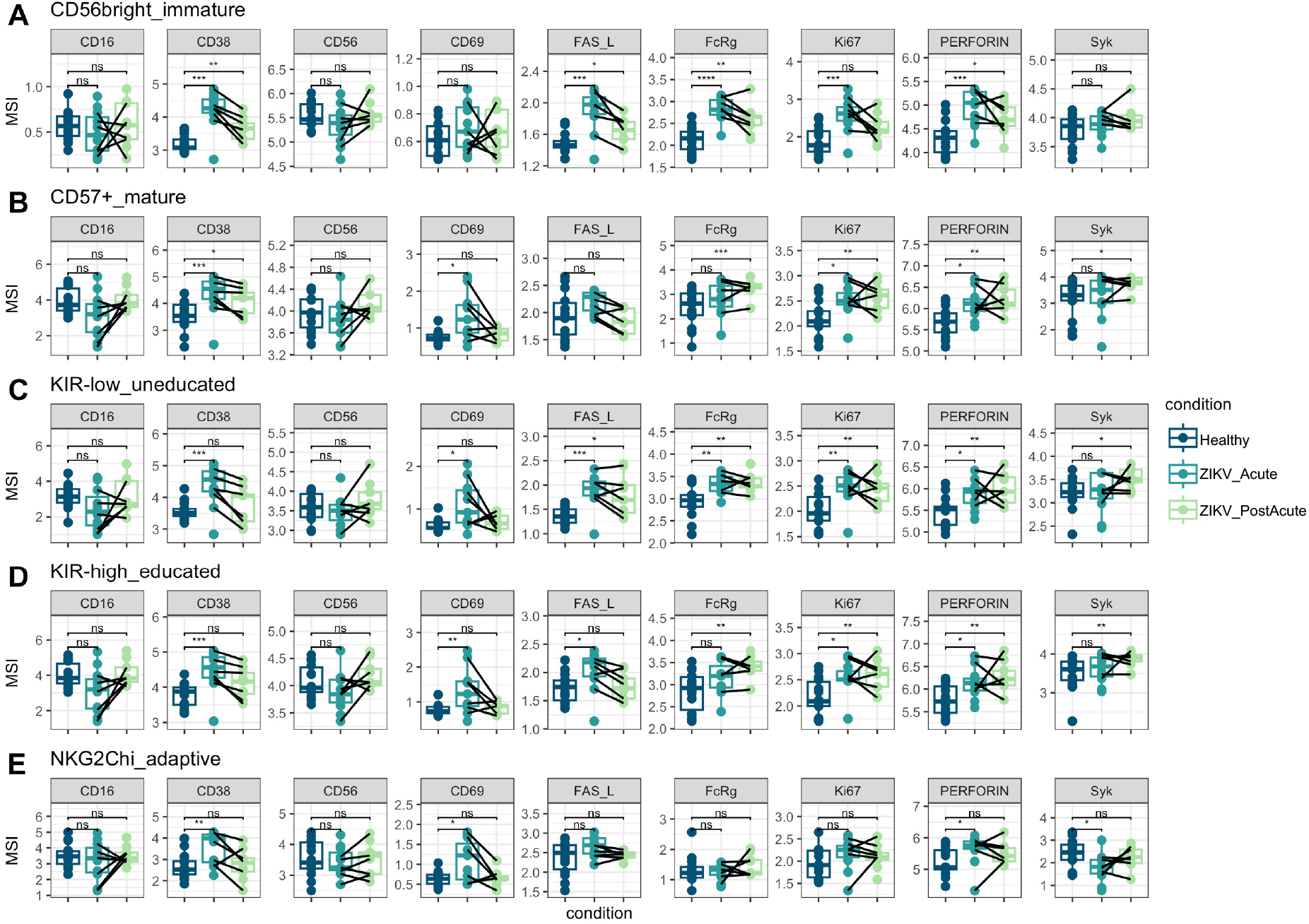
MSI of selected NK markers by maturation & education status a) Immature (CD56^bright^CD16^-^, cluster 8) b) Mature (CD57^+^, clusters 1, 2, and 6) c) Uneducated (KIR^low^, clusters 3, 7, and 8) d) Educated (KIR^high^, clusters 1, 2, 6, and 9) e) Adaptive (NKG2C^high^, cluster 1)

**Fig S4:**
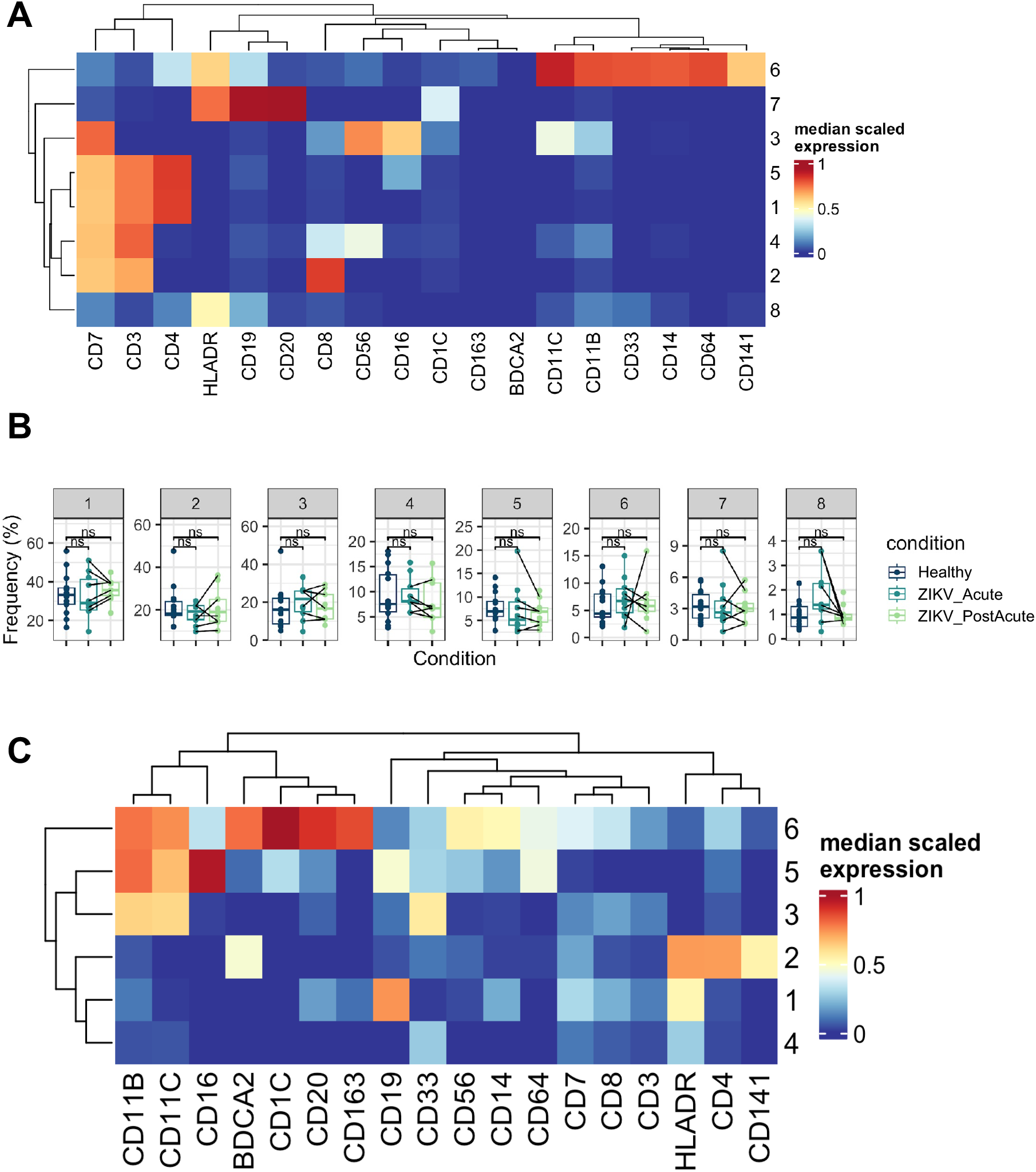
Unsupervised clustering of PBMCs a) Heatmap of median protein expression and b) Cluster frequencies of original parc clustering (resolution = 0.12) before merging of cluster 1 and 5 to CD4 T cells and clusters 2 and 4 for CD8 T cell clusters c) Re-clustering of cells in original cluster 8 with resolution 0.12

**Fig S5:**
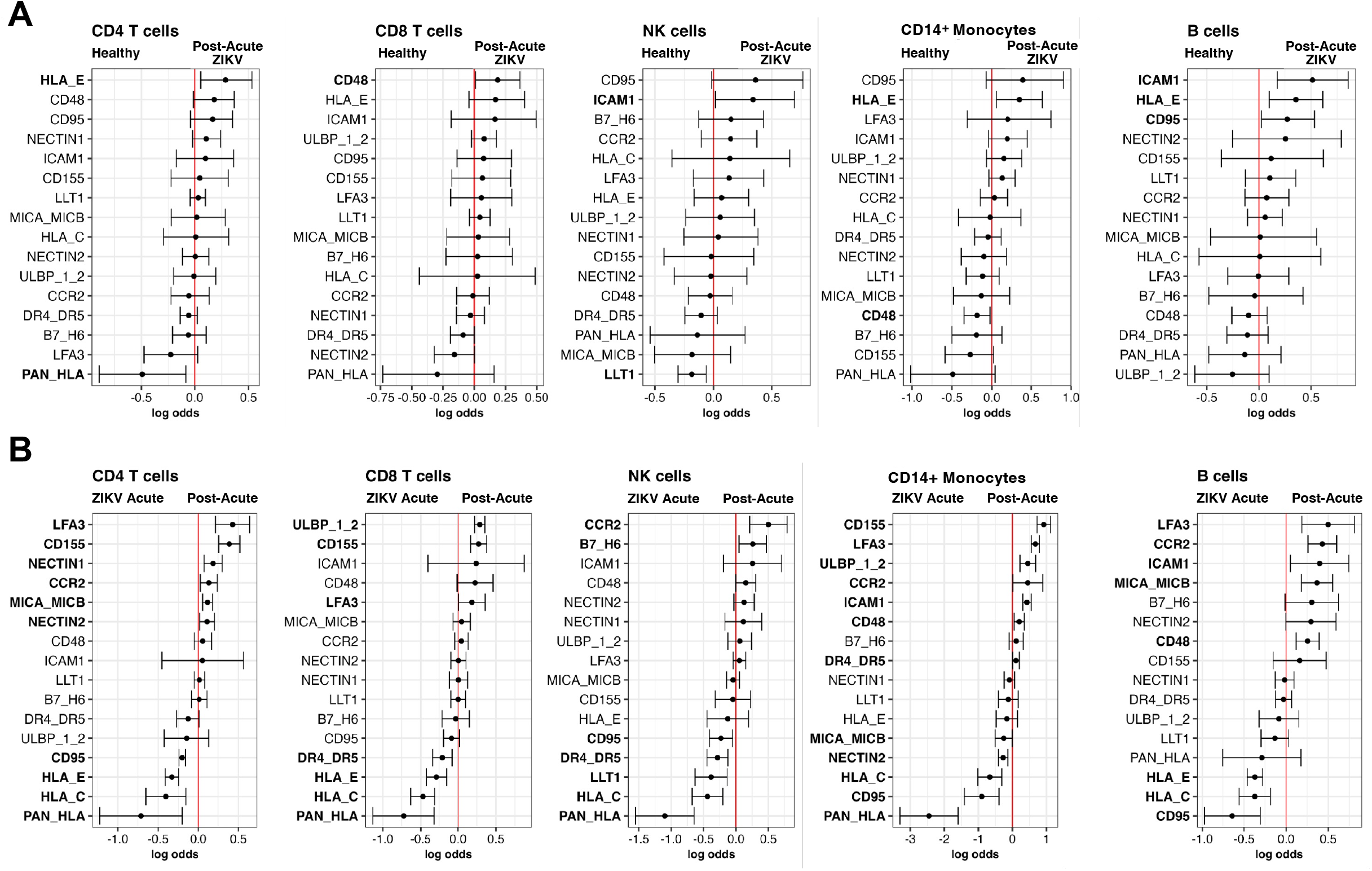
CytoGLM of PBMC subsets a) Healthy (n = 13-14) vs. Post-acute (n = 9-10) b) Acute vs. Post-acute (n = 7 pairs)

**Fig S6:**
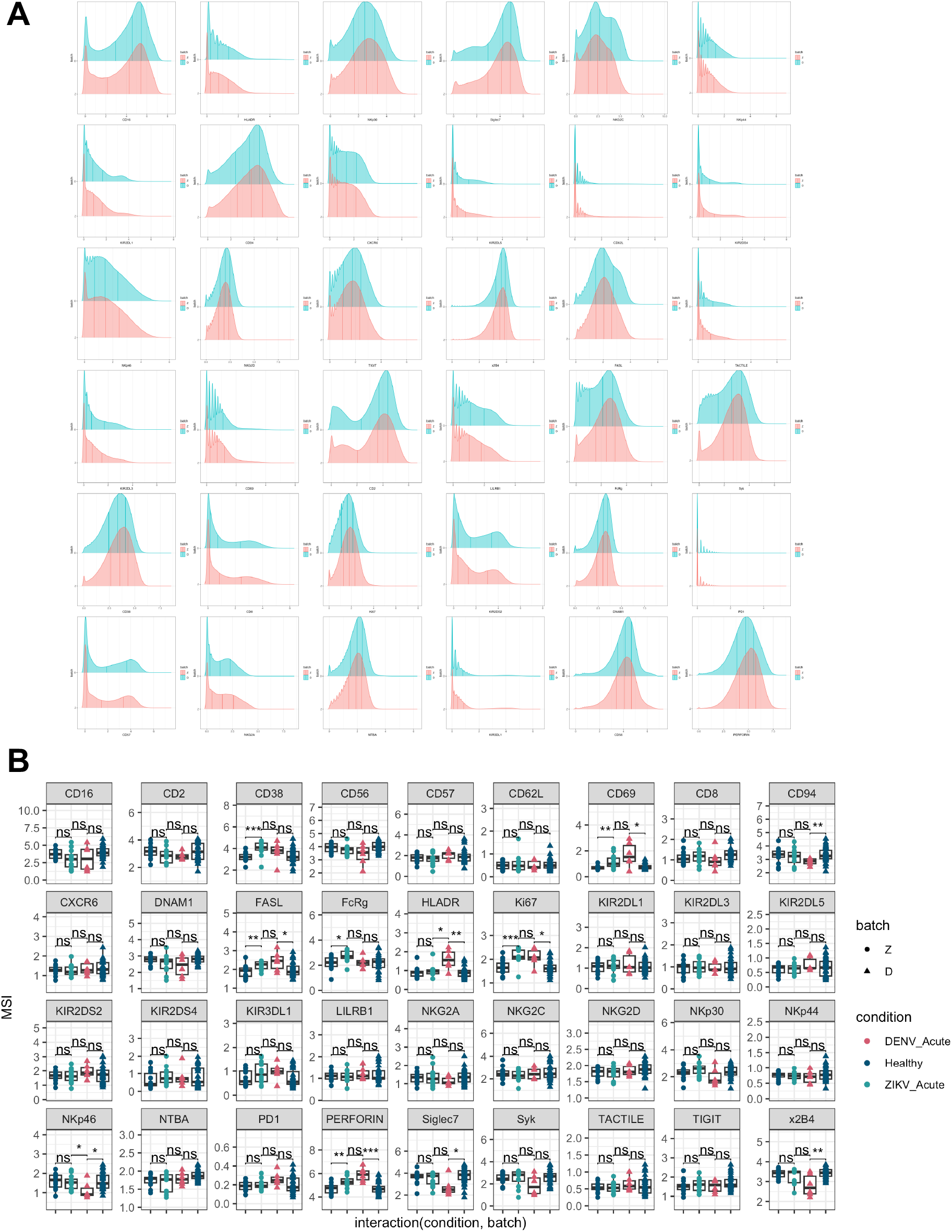
Integration of ZIKV and DENV CyTOF datasets. a) Marker distributions by batch (Z=this study, D=McKechnie *et al* 2020 [20]) b) Mean signal intensity (MSI) of all NK cell markers after data integration

**Table S1:** Ligand CyTOF panel info

**Table S2:** NK CyTOF panel info

**File S1:** Differential state testing results of Acute vs. Post-acute and Post-acute vs. Healthy in PBMC clusters

**Table S1:** NK Ligand CyTOF panel. Antibodies in red were spiked into the frozen panel separately

| Isotope | Protein | Vendor | Clone | Note |
| --- | --- | --- | --- | --- |
| 89Y | HLA-DR | Biolegend | L243 | surface |
| 112Cd | CD19 Qdot | Life Technologies | SJ25-C1 | surface |
| 115In | CD3 | Biolegend | UCHT1 | surface |
| 141Pr | CD20 | Biolegend | 2H7 | surface |
| 142Nd | CD163 | Biolegend | GHI/61 | surface |
| 143Nd | PAN HLA | Biolegend | W6/32 | surface |
| 144Nd | CD7 | Biolegend | CD7-6B7 | surface |
| 145Nd | CD8 | Biolegend | SK1 | surface |
| 146Nd | CD48 | Biolegend | BJ40 | surface |
| 147Sm | BDCA-2 | Biolegend | 201A | surface |
| 148Nd | ICAM1 (CD54) | Biolegend | HA58 | surface |
| 149Sm | LLT-1 | R&D | 402659 | surface |
| 151Eu | CD4 | Biolegend | OKT4 | surface |
| 152Sm | CD64 | Biolegend | 10.1 | surface |
| 153Eu | HLA-C | Millipore | DT9 | surface |
| 154Sm | CCR2 | Biolegend | K036C2 | surface |
| 155Gd | HLA-E | Biolegend | 3D12 | surface |
| 156Gd | CD95 | Biolegend | DX2 | surface |
| 157Gd | Nectin-1 (CD111) | Biolegend | R1.302 | surface |
| 158Gd | MICA/MICB | R&D | 159227/236511 | surface |
| 159Tb | DR4 (CD261) | Biolegend | DJR1 | surface |
| 159Tb | DR5 (CD262) | Biolegend | DJR2-2 | surface |
| 160Gd | CD1c | Biolegend | L161 | surface |
| 161Dy | ULBP-1/2 (5 and 6) | R&D | 170818/165903 | surface |
| 162Dy | CD11c | Biolegend | Bu15 | surface |
| 164Dy | Nectin-2 (CD112) | Biolegend | TX31 | surface |
| 165Ho | CD155 (PVR) | Biolegend | SKII.4 | surface |
| 166Er | HLA-Bw4 | Miltenyi | REA274 | surface |
| 167Er | CD32 | STEMCELL | IV.3 | surface |
| 168Er | HLA-Bw6 | Miltenyi | REA143/ 130-095-212 | surface |
| 169Tm | CD14 | Biolegend | M5E2 | surface |
| 170Er | CD11b | Biolegend | ICRF44 | surface |
| 171Yb | LFA-3 (CD58) | Biolegend | TS2/9 | surface |
| 172Yb | CD33 | Biolegend | WM53 | surface |
| 173Yb | CD141 | Fluidigm pre-conjugated | 1A4 | surface |
| 174Yb | CD56 | BD Pharmingen | NCAM16.2 | surface |
| 175Lu | CD86 | Biolegend | IT2.2 | surface |
| 176Yb | B7-H6 | R&D | 875001 | surface |
| 209Bi | CD16 | Fluidigm pre-conjugated | 3G8 | surface |
| 150Nd | Flavi E-protein | EMD Millipore | D1-4G2-4-15 | intracellular |

**Table S2:**
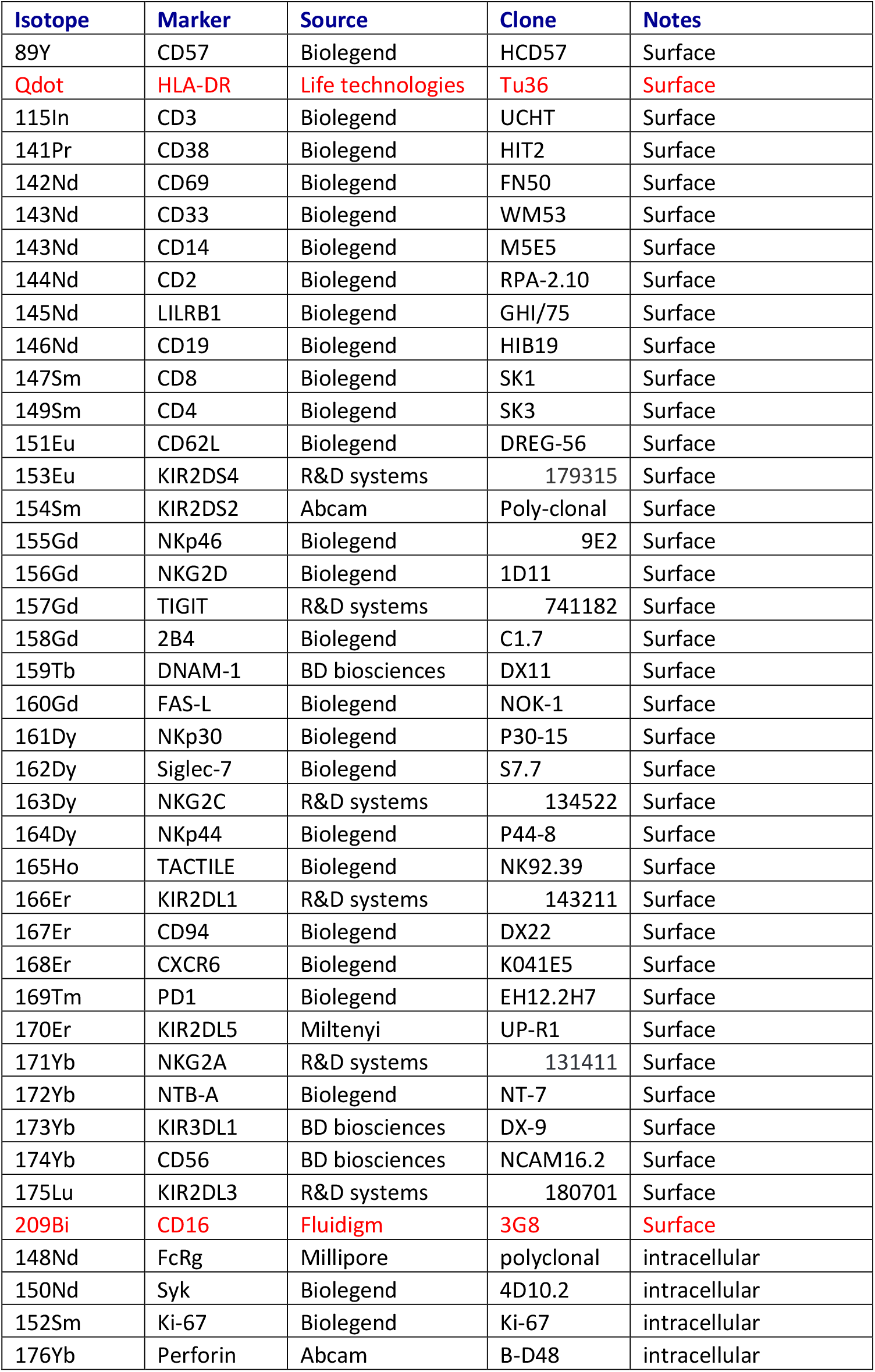
NK cell CyTOF panel. Antibodies in red were spiked into the lyophilized panel separately.

| Isotope | Marker | Source | Clone | Notes |
| --- | --- | --- | --- | --- |
| 89Y | CD57 | Biolegend | HCD57 | Surface |
| Qdot | HLA-DR | Life technologies | Tu36 | Surface |
| 115In | CD3 | Biolegend | UCHT | Surface |
| 141Pr | CD38 | Biolegend | HIT2 | Surface |
| 142Nd | CD69 | Biolegend | FN50 | Surface |
| 143Nd | CD33 | Biolegend | WM53 | Surface |
| 143Nd | CD14 | Biolegend | M5E5 | Surface |
| 144Nd | CD2 | Biolegend | RPA-2.10 | Surface |
| 145Nd | LILRB1 | Biolegend | GHI/75 | Surface |
| 146Nd | CD19 | Biolegend | HIB19 | Surface |
| 147Sm | CD8 | Biolegend | SK1 | Surface |
| 149Sm | CD4 | Biolegend | SK3 | Surface |
| 151Eu | CD62L | Biolegend | DREG-56 | Surface |
| 153Eu | KIR2DS4 | R&D systems | 179315 | Surface |
| 154Sm | KIR2DS2 | Abcam | Poly-clonal | Surface |
| 155Gd | NKp46 | Biolegend | 9E2 | Surface |
| 156Gd | NKG2D | Biolegend | 1D11 | Surface |
| 157Gd | TIGIT | R&D systems | 741182 | Surface |
| 158Gd | 2B4 | Biolegend | C1.7 | Surface |
| 159Tb | DNAM-1 | BD biosciences | DX11 | Surface |
| 160Gd | FAS-L | Biolegend | NOK-1 | Surface |
| 161Dy | NKp30 | Biolegend | P30-15 | Surface |
| 162Dy | Siglec-7 | Biolegend | S7.7 | Surface |
| 163Dy | NKG2C | R&D systems | 134522 | Surface |
| 164Dy | NKp44 | Biolegend | P44-8 | Surface |
| 165Ho | TACTILE | Biolegend | NK92.39 | Surface |
| 166Er | KIR2DL1 | R&D systems | 143211 | Surface |
| 167Er | CD94 | Biolegend | DX22 | Surface |
| 168Er | CXCR6 | Biolegend | K041E5 | Surface |
| 169Tm | PD1 | Biolegend | EH12.2H7 | Surface |
| 170Er | KIR2DL5 | Miltenyi | UP-R1 | Surface |
| 171Yb | NKG2A | R&D systems | 131411 | Surface |
| 172Yb | NTB-A | Biolegend | NT-7 | Surface |
| 173Yb | KIR3DL1 | BD biosciences | DX-9 | Surface |
| 174Yb | CD56 | BD biosciences | NCAM16.2 | Surface |
| 175Lu | KIR2DL3 | R&D systems | 180701 | Surface |
| 209Bi | CD16 | Fluidigm | 3G8 | Surface |
| 148Nd | FcRg | Millipore | polyclonal | intracellular |
| 150Nd | Syk | Biolegend | 4D10.2 | intracellular |
| 152Sm | Ki-67 | Biolegend | Ki-67 | intracellular |
| 176Yb | Perforin | Abcam | B-D48 | intracellular |

## References

1. Bhatt S, Gething PW, Brady OJ, Messina JP, Farlow AW, Moyes CL, et al. The global distribution and burden of dengue. Nature. 2013;496: 504–507.

2. van Leur SW, Heunis T, Munnur D, Sanyal S. Pathogenesis and virulence of flavivirus infections. Virulence. 2021;12: 2814–2838.

3. Lanciotti RS, Kosoy OL, Laven JJ, Velez JO, Lambert AJ, Johnson AJ, et al. Genetic and serologic properties of Zika virus associated with an epidemic, Yap State, Micronesia, 2007. Emerg Infect Dis. 2008;14: 1232–1239.

4. Wilk AJ, Blish CA. Diversification of human NK cells: Lessons from deep profiling. J Leukoc Biol. 2018;103: 629–641.

5. Blish CA. Natural killer cell diversity in viral infection: Why and how much? Pathog Immun. 2016;1: 165–192.

6. Horowitz A, Strauss-Albee DM, Leipold M, Kubo J, Nemat-Gorgani N, Dogan OC, et al. Genetic and environmental determinants of human NK cell diversity revealed by mass cytometry. Sci Transl Med. 2013;5: 208ra145.

7. Strauss-Albee DM, Fukuyama J, Liang EC, Yao Y, Jarrell JA, Drake AL, et al. Human NK cell repertoire diversity reflects immune experience and correlates with viral susceptibility. Sci Transl Med. 2015;7: 297ra115.

8. Marquardt N, Ivarsson MA, Blom K, Gonzalez VD, Braun M, Falconer K, et al. The human NK cell response to yellow fever virus 17D is primarily governed by NK cell differentiation independently of NK cell education. J Immunol. 2015;195: 3262–3272.

9. Blom K, Braun M, Pakalniene J, Lunemann S, Enqvist M, Dailidyte L, et al. NK cell responses to human tick-borne encephalitis virus infection. J Immunol. 2016;197: 2762–2771.

10. Yao Y, Strauss-Albee DM, Zhou JQ, Malawista A, Garcia MN, Murray KO, et al. The natural killer cell response to West Nile virus in young and old individuals with or without a prior history of infection. PLoS One. 2017;12: e0172625.

11. Ghita L, Yao Z, Xie Y, Duran V, Cagirici HB, Samir J, et al. Global and cell type-specific immunological hallmarks of severe dengue progression identified via a systems immunology approach. Nat Immunol. 2023;24: 2150–2163.

12. Robinson ML, Glass DR, Duran V, Agudelo Rojas OL, Sanz AM, Consuegra M, et al. Magnitude and kinetics of the human immune cell response associated with severe dengue progression by single-cell proteomics. Sci Adv. 2023;9: eade7702.

13. Azeredo EL, De Oliveira-Pinto LM, Zagne SM, Cerqueira DIS, Nogueira RMR, Kubelka CF. NK cells, displaying early activation, cytotoxicity and adhesion molecules, are associated with mild dengue disease. Clin Exp Immunol. 2006;143: 345–356.

14. Maucourant C, Nonato Queiroz GA, Corneau A, Leandro Gois L, Meghraoui-Kheddar A, Tarantino N, et al. NK Cell Responses in Zika Virus Infection Are Biased towards Cytokine-Mediated Effector Functions. J Immunol. 2021;207: 1333–1343.

15. McKechnie JL, Beltrán D, Pitti A, Saenz L, Araúz AB, Vergara R, et al. HLA Upregulation During Dengue Virus Infection Suppresses the Natural Killer Cell Response. Front Cell Infect Microbiol. 2019;9: 268.

16. Lanciotti RS, Calisher CH, Gubler DJ, Chang GJ, Vorndam AV. Rapid detection and typing of dengue viruses from clinical samples by using reverse transcriptase-polymerase chain reaction. J Clin Microbiol. 1992;30: 545–551.

17. Lanciotti RS, Kosoy OL, Laven JJ, Panella AJ, Velez JO, Lambert AJ, et al. Chikungunya virus in US travelers returning from India, 2006. Emerg Infect Dis. 2007;13: 764–767.

18. Vendrame E, McKechnie JL, Ranganath T, Zhao NQ, Rustagi A, Vergara R, et al. Profiling of the Human Natural Killer Cell Receptor-Ligand Repertoire. J Vis Exp. 2020. doi:10.3791/61912

19. Hamlin RE, Pienkos SM, Chan L, Stabile MA, Pinedo K, Rao M, et al. Sex differences and immune correlates of Long Covid development, symptom persistence, and resolution. Sci Transl Med. 2024;16: eadr1032.

20. McKechnie JL, Beltrán D, Ferreira A-MM, Vergara R, Saenz L, Vergara O, et al. Mass Cytometry Analysis of the NK Cell Receptor-Ligand Repertoire Reveals Unique Differences between Dengue-Infected Children and Adults. Immunohorizons. 2020;4: 634–647.

21. Gherardini PF. premessa: R package for pre-processing of mass and flow cytometry data. Github; 2023. Available: https://github.com/ParkerICI/premessa

22. Ellis B, Haaland P, Hahne F, Le Meur N, Gopalakrishnan N, Spidlen J, et al. flowCore: flowCore: Basic structures for flow cytometry data. 2024. doi:10.18129/B9.bioc.flowCore

23. Crowell HL, Zanotelli VRT, Chevrier S, Robinson MD. CATALYST: Cytometry dATa anALYSis Tools. 2024. doi:10.18129/B9.bioc.CATALYST

24. Stassen SV, Siu DMD, Lee KCM, Ho JWK, So HKH, Tsia KK. PARC: ultrafast and accurate clustering of phenotypic data of millions of single cells. Bioinformatics. 2020;36: 2778–2786.

25. Weber LM, Nowicka M, Soneson C, Robinson MD. diffcyt: Differential discovery in highdimensional cytometry via high-resolution clustering. Commun Biol. 2019;2: 183.

26. Seiler C, Ferreira A-M, Kronstad LM, Simpson LJ, Le Gars M, Vendrame E, et al. CytoGLMM: conditional differential analysis for flow and mass cytometry experiments. BMC Bioinformatics. 2021;22: 137.

27. Pedersen CB, Dam SH, Barnkob MB, Leipold MD, Purroy N, Rassenti LZ, et al. cyCombine allows for robust integration of single-cell cytometry datasets within and across technologies. Nat Commun. 2022;13: 1698.

28. Lanciotti RS, Lambert AJ, Holodniy M, Saavedra S, Signor LDCC. Phylogeny of Zika virus in western hemisphere, 2015. Emerg Infect Dis. 2016;22: 933–935.

29. Baz M. Zika virus isolation, purification, and titration. Methods Mol Biol. 2020;2142: 9–22.

30. Gu Z, Eils R, Schlesner M. Complex heatmaps reveal patterns and correlations in multidimensional genomic data. Bioinformatics. 2016;32: 2847–2849.

31. Wickham H. ggplot2: Elegant Graphics for Data Analysis. Springer-Verlag New York; 2016. Available: https://ggplot2.tidyverse.org

32. Kassambara A. _ggpubr: “ggplot2” Based Publication Ready Plots. Comprehensive R Archive Network (CRAN); 2025. Available: https://CRAN.R-project.org/package=ggpubr

33. Liu W, Scott JM, Langguth E, Chang H, Park PH, Kim S. FcRγ gene editing reprograms conventional NK cells to display key features of adaptive human NK cells. iScience. 2020;23: 101709.

34. McDonald EM, Anderson J, Wilusz J, Ebel GD, Brault AC. Zika virus replication in myeloid cells during acute infection is vital to viral dissemination and pathogenesis in a mouse model. J Virol. 2020;94. doi:10.1128/JVI.00838-20

35. Santara SS, Crespo ÂC, Mulik S, Ovies C, Boulenouar S, Strominger JL, et al. Decidual NK cells kill Zika virus–infected trophoblasts. Proceedings of the National Academy of Sciences. 2021;118: e2115410118.

36. Glasner A, Oiknine-Djian E, Weisblum Y, Diab M, Panet A, Wolf DG, et al. Zika Virus Escapes NK Cell Detection by Upregulating Major Histocompatibility Complex Class I Molecules. J Virol. 2017;91. doi:10.1128/JVI.00785-17

37. Barnard TR, Rajah MM, Sagan SM. Contemporary Zika Virus Isolates Induce More dsRNA and Produce More Negative-Strand Intermediate in Human Astrocytoma Cells. Viruses. 2018;10. doi:10.3390/v10120728

38. Zimmer CL, Cornillet M, Solà-Riera C, Cheung K-W, Ivarsson MA, Lim MQ, et al. NK cells are activated and primed for skin-homing during acute dengue virus infection in humans. Nat Commun. 2019;10: 3897.

39. Barros JB de S, da Silva PAN, Koga R de CR, Gonzalez-Dias P, Carmo Filho JR, Nagib PRA, et al. Acute Zika virus infection in an endemic area shows modest proinflammatory systemic immunoactivation and cytokine-symptom associations. Front Immunol. 2018;9: 821.

40. Lee J, Zhang T, Hwang I, Kim A, Nitschke L, Kim M, et al. Epigenetic modification and antibody-dependent expansion of memory-like NK cells in human cytomegalovirus-infected individuals. Immunity. 2015;42: 431–442.

41. Shemesh A, Su Y, Calabrese DR, Chen D, Arakawa-Hoyt J, Roybal KT, et al. Diminished cell proliferation promotes natural killer cell adaptive-like phenotype by limiting FcεRIγ expression. J Exp Med. 2022;219. doi:10.1084/jem.20220551

42. Tappe D, Pérez-Girón JV, Zammarchi L, Rissland J, Ferreira DF, Jaenisch T, et al. Cytokine kinetics of Zika virus-infected patients from acute to reconvalescent phase. Med Microbiol Immunol. 2016;205: 269–273.

43. Lum F-M, Lye DCB, Tan JJL, Lee B, Chia P-Y, Chua T-K, et al. Longitudinal study of cellular and systemic cytokine signatures to define the dynamics of a balanced immune environment during disease manifestation in Zika virus-infected patients. J Infect Dis. 2018;218: 814–824.

44. Lopez-Vergès S, Milush JM, Pandey S, York VA, Arakawa-Hoyt J, Pircher H, et al. CD57 defines a functionally distinct population of mature NK cells in the human CD56dimCD16+ NK-cell subset. Blood. 2010;116: 3865–3874.

45. Sen Santara S, Lee D-J, Crespo Â, Hu JJ, Walker C, Ma X, et al. The NK cell receptor NKp46 recognizes ecto-calreticulin on ER-stressed cells. Nature. 2023;616: 348–356.

46. Drews E, Adam A, Htoo P, Townsley E, Mathew A. Upregulation of HLA-E by dengue and not Zika viruses. Clin Transl Immunology. 2018;7: e1039.

47. Airo AM, Urbanowski MD, Lopez-Orozco J, You JH, Skene-Arnold TD, Holmes C, et al. Expression of flavivirus capsids enhance the cellular environment for viral replication by activating Akt-signalling pathways. Virology. 2018;516: 147–157.

48. Lee JK, Kim J-A, Oh S-J, Lee E-W, Shin OS. Zika virus induces tumor necrosis factor-related apoptosis inducing ligand (TRAIL)-mediated apoptosis in human neural progenitor cells. Cells. 2020;9: 2487.

49. Maucourant C, Queiroz GAN, Samri A, Grassi MFR, Yssel H, Vieillard V. Zika virus in the eye of the cytokine storm. Eur Cytokine Netw. 2019;30: 74–81.

